# BDKANN - Biological Domain Knowledge-based Artificial Neural Network for drug response prediction

**DOI:** 10.1101/840553

**Authors:** Oliver Snow, Hossein Sharifi-Noghabi, Jialin Lu, Olga Zolotareva, Mark Lee, Martin Ester

## Abstract

**Motivation:** One of the main goals of precision oncology is to predict the response of a patient to a given cancer treatment based on their genomic profile. Although current models for drug response prediction are becoming more accurate, they are also ‘black boxes’ and cannot explain their predictions, which is of particular importance in cancer treatment. Many models also do not leverage prior biological knowledge, such as the hierarchical information on how proteins form complexes and act together in pathways.

**Results:** In this work, we use this prior biological knowledge to form the architecture of a deep neural network to predict cancer drug response from cell line gene expression data. We find that our approach not only has a low prediction error compared to baseline models but also allows meaningful interpretation of the network. These interpretations can both explain predictions made and discover novel connections in the biological knowledge that may lead to new hypotheses about mechanisms of drug action.

**Availability:** Code at https://github.com/osnow/BDKANN

**Supplementary information:** Included with submission

## 1 Introduction

A major goal of precision oncology is to provide cancer patients with the most effective therapies given their underlying genomic profile. One of the major challenges to achieve this goal is to build an accurate computational model to predict drug response given the omics data of apatient (Ali and Aittokallio, 2019; Marquart *et al*., 2018). Unfortunately, current clinical datasets with drug response labels are not large enough to train an accurate model while gathering more data is expensive and time consuming. In addition, it is impossible to treat a patient with multiple drugs independently to separately study their effects. However, numerous pre-clinical datasets have recently been created based on human cancer cell lines and patient-derived xenografts (Iorio *et al*., 2016; Gao *et al*., 2015; Garnett *et al*., 2012; Barretina *et al*., 2012). These pre-clinical datasets are larger than clinical datasets and are screened with tens or hundreds of drugs which makes them useful resources for training a computational model.

For drug response prediction on pre-clinical datasets, previous studies have proposed different methods such as regression (Geeleher *et al*., 2014), kernel learning (He *et al*., 2018), deep neural networks (DNNs) (Sharifi-Noghabi *et al*., 2019; Ding *et al*., 2018), and transfer learning (Noghabi *et al*., 2020; Mourragui *et al*., 2019) to map the given omics data–often gene expression–to a measure of response such as the IC50 or area above the dose-response curve (AAC) (Smirnov *et al*., 2017). Although these methods have shown promising results, they have not considered prior biological knowledge in their models. These methods used genes as the features of their models in isolation, however, genes do not operate in isolation, but rather work in biological networks or pathways. In the context of drug response prediction, genes and the corresponding proteins work together to form protein complexes, which then operate in different pathways, and eventually the function of a pathway is disrupted by a drug. Therefore, our hypothesis is that incorporating this domain expert knowledge in a computational model should improve the accuracy of drug response prediction by avoiding over-fitting on a rather small dataset.

The other major challenge is that despite the relatively accurate predictions of existing models, most are so called ‘black boxes’ in which the reasoning behind the prediction is unknown. This is particularly undesirable in the context of cancer therapy as clinicians require a rational explanation of why a drug is expected to work for a given patient, however, current models do not provide this explanation. This motivates the use of domain expert knowledge as the structure of the model so that predictions can then be traced back to specific nodes in the network that represent real biological entities such as pathways, complexes, or specific genes. Ma *et al*. (2018) was one of the first attempts at creating an interpretable deep neural network which predicted yeast growth using a model made from a network of gene ontology (GO) terms. This model, called DCELL, was able to accurately predict cell growth from genotype data (mutation pairs) while also being able to explain what components of the cell were involved in the prediction.

The work by Gaudelet *et al*. (2019) also used prior domain knowledge to define the structure of a neural network for the task of disease diagnosis based on gene expression data. Rather than using a GO network, the proposed method uses biological pathway and protein complex data to define the structure of the neural network. The method achieved accuracy comparable to that of other methods and was also able to uncover the underlying biological entities responsible for a diagnosis and to discover disease-disease relationships.

Similarly, Hao *et al*. (2018) built a DNN model called PASNet that uses biological pathway (but not protein complex) information to construct the connections in a neural network to predict survival of cancer patients. The proposed model outperformed other predictors of patient survival while also being able to identify key pathways involved in progression of the disease.

Lastly, in another study, Kang *et al*. (2017) constructed a neural network based on gene regulatory network knowledge to predict the response to different therapies. A significant contribution of this work was to regularize those connections in the network not supported by the gene regulatory information using L1 regularization. This effectively biases the network towards the connections defined by the prior biological knowledge while still allowing the possibility for new connections in the network to remain.

Together, these previous works show the potential of prior biological knowledge for defining the architecture of DNNs, both for improving prediction accuracy and for creating interpretable models with functional explanations for the predictions made. Drawing on the strengths of these past works we build an interpretable DNN based on biological knowledge for the task of drug response prediction which we call BDKANN. We also make a extended version of BDKANN that we call BDKANN+, which has the added ability to discover new connections in the biological domain knowledge through the use of regularization. We compare both versions of our model to both knowledge-based and non knowledge-based DNN baselines and find that not only does BDKANN+ outperform BDKANN and baselines but also allows for meaningful interpretation of those results that can help us better understand the decisions of the model and help generate hypotheses relevant to cancer drug response prediction.

## 2 Approach

In this paper, we employ prior biological knowledge in the form of the hierarchical connections of genes to protein complexes to pathways and finally to drugs as layers of a DNN. The structure of BDKANN (and BDKANN+) is as follows: 1) in the gene layer of this network, a node represents a gene for which the expression data is available. 2) In the protein complex layer, a node represents the complex that genes in the previous layer can form. 3) In the pathway layer, a node represents a pathway that a protein complex (or multiple complexes) in the previous layer is (are) a part of. Lastly, 4) in the drug layer, a node represents a drug that targets a given pathway(s) in the previous layer.

We connect these layers in the neural network using the biological knowledge obtained from Reactome, an open source, peer-reviewed, and manually curated database containing pathway, complex, and even drugtarget information (Fabregat *et al*., 2017). The advantage of this approach is that the DNN model predicts the response to multiple drugs using prior biological knowledge. However, it cannot discover new associations between different layers of information in the context of drug response. Therefore, we add edges to make the network fully connected but also add regularization on these additional edges, which are not supported by the biological knowledge, to shrink their weights towards zero. Thus, the proposed method exploits the biological knowledge and is also capable of discovering new associations if an edge which is not supported by the biological knowledge “survives” the regularization.

## 3 Methods

### 3.1 Problem definition

Given gene expression data *X* ∈ *M* × *N*, where *M* is the number of cell lines and *N* is the number of genes, and area above the dose-response curve (AAC) labels *Y* ∈ *M* × *K*, where *K* is the number of screened drugs, the objective is to learn a model *f*(*X*) to predict *Y*. In this paper, *f* is a DNN parameterized by Θ that receives gene expression data from cell lines and predicts the response to *K* drugs such that it minimizes the total Mean Squared Error. The goal is to incorporate the biological knowledge in designing the structure and connections of different layers in the DNN implementing *f*, and at the same time explore new connections and associations between different layers beyond the available domain expert knowledge. For example, for any two consecutive layers such as *L_i_* and *L_j_*, the connections (weights of the neural network) are 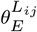 and 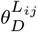 such that 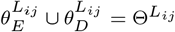, where 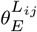 are connections supported by the domain expert knowledge, and 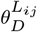 are the additional connections (i.e. unknown based on available domain expert knowledge) for discovering new associations between *L_i_* and *L_j_*. To avoid overparameterization of *f*, 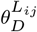 will be regularized such that only connections with direct impact on the predictions survive by forcing the others without any impact to become zero.

### 3.2 Datasets

In this work, we utilized two datasets: 1) The Genomics of Drug Sensitivity in Cancer (GDSC) dataset (Iorio *et al*., 2016), consisting of more than 1000 pan-cancer cell lines, screened with 265 targeted and chemotherapy drugs. 2) The Cancer Therapeutic Response Portal (CTRPv2) dataset (Rees *et al*., 2016), consisting of more than 800 pan-cancer cell lines, screened with 481 targeted and chemotherapy drugs. The general make-up of these two datasets can be seen in table 1, which shows the number of genes, drugs, and cell lines tested in each dataset as well as the overlap of these entities and the correlation of the response variable (AAC). Despite each dataset containing many cell lines and drugs, only 92 drugs and 592 cell lines are covered by both datasets, and within that overlap there are still many missing values for drug-cell line pairs. Additionally, the correlation between the response variables of the two datasets is fairly poor and we chose to use AAC over IC50 due to its higher correlation. There has been some discussion on why the measured drug response differs between these two datasets and the problems that poses for training machine learning models on them (Consortium *et al*., 2015; Safikhani *et al*., 2016). We recognize that these datasets have distinct distributions and evaluate our model’s ability to generalize from one to the other.

**Table 1.** Descriptive statistics comparing GDSC and CTRPv2 datasets. *Values calculated for only the drugs and cell lines that overlapped between the two datasets

|  | GDSC | CTRPv2 |
| --- | --- | --- |
| # of cell lines | 1109 | 888 |
| # of drugs | 250 | 544 |
| # of genes | 18645 | 20342 |
| Average AAC* | 0.18 +/- 0.14 | 0.21 +/- 0.13 |
| AAC missing* | 1406 | 495 |
| AAC corr. | 0.503 |  |
| IC50 corr. | 0.466 |  |
| Shared drugs | 92 |  |
| Shared cell lines | 592 |  |

We obtained the GDSC and CTRPv2 datasets via PharmacoGx R package (Smirnov *et al*., 2015) and PharmacoDB (Smirnov *et al*., 2017). These two datasets have 93 drugs in common, however, we focused on 11 of those drugs (Doxorubicin, Tamoxifen, Masitinib, 17-AAG, GDC-0941, ATRA, Gefitinib, BIBW2992, MK-2206, AZD6244, and PLX4720). We chose to only include drugs with sufficient pathway information available in Reactome and we filtered out drugs with a high fraction of NA values for the drug response outcome. Our model could easily be extended to predict for more drugs when response data is collected on more cell lines and more complete pathway-drug information is acquired.

### 3.3 BDKANN architecture

The architecture of BDKANN is defined by the gene expression and drug response data and the acquired biological knowledge from the network data in Reactome. The general architecture can be seen in figure 1, where the input layer consists of genes, the first hidden layer consists of protein complexes, the second hidden layer consists of pathways, and the output layer corresponds to the drugs for which we predict AAC. The dimension of the input layer is 607, defined by the set of landmark genes whose expression was measured in both studied datasets and reduced to the genes for which there is complex information available. Landmark genes are those defined by Peck et al. (2006), which have been shown to be predictive of the expression of the remaining 85% of genes (Subramanian *et al*., 2017). We reduced the input to only those landmark genes in order to have the same input dimension for all the models we tested and the DCELL method ran into significant memory issues when run with all available genes. The dimension of the first hidden layer is 4448, corresponding to the number of protein complexes that the input genes map to. The dimension of the second hidden layer is 867, corresponding to the number of pathways that the previous complexes map to. The final output layer dimension is 11, corresponding to the number of overlapping drugs between the datasets and reduced those drugs with drug response data for more than 50% of cell lines.

**Fig. 1:**
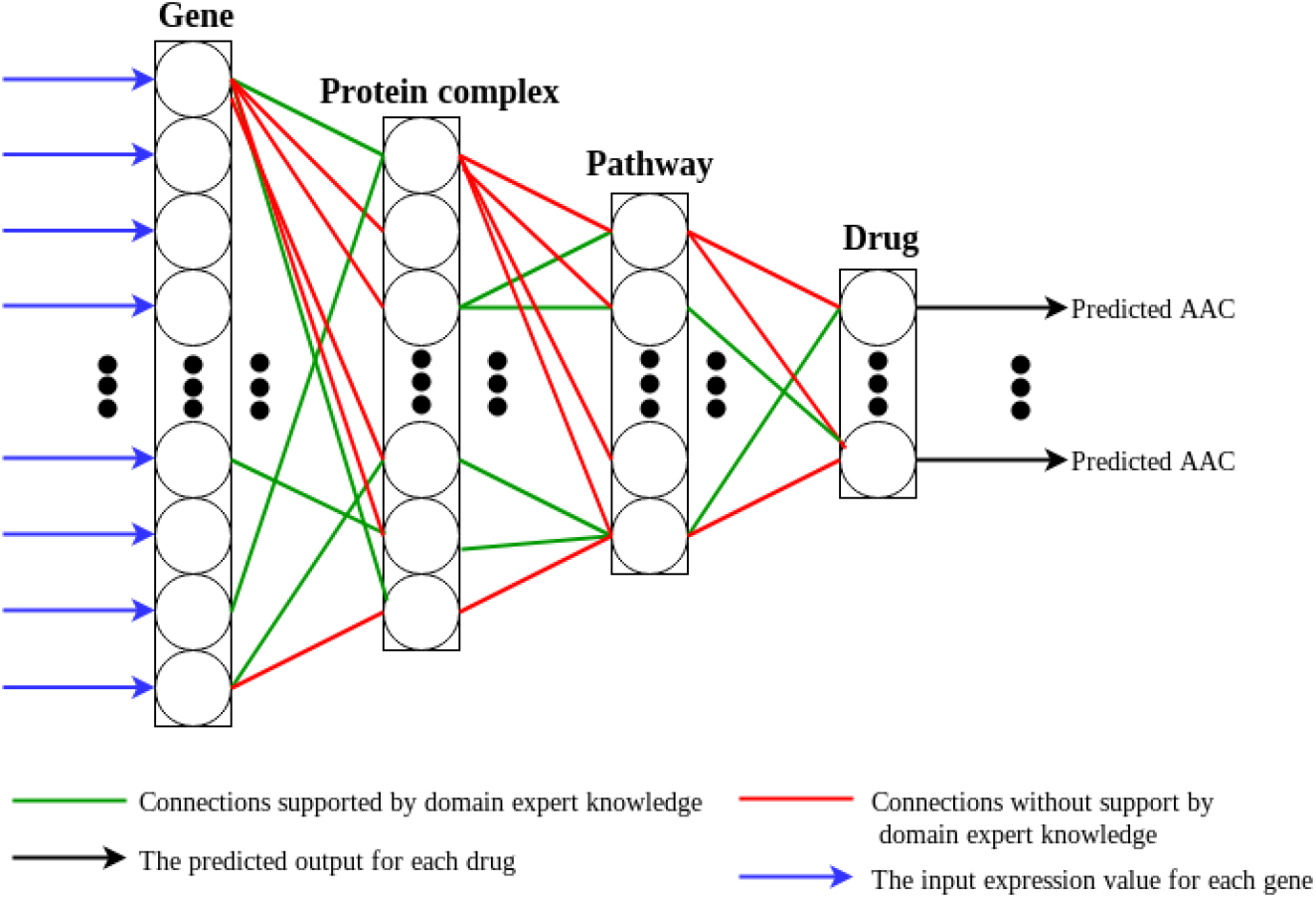
Overview of the structure of BDKANN. BDKANN+ has added connections (red) that are not supported by domain knowledge but are regularized to give preference to the green connections (supported by biological knowledge)

### 3.4 Cost function

A key part of BDKANN+ is the discovery regularization. The goal of this regularization is to discover new associations between different layers of the network. We employ *l*_1_ regularization to force the weights corresponding to the edges that are not supported by the biological domain knowledge to become zero. The rationale behind regularizing these edges is that we want to give preference to the prior biological knowledge, but we also recognize that this knowledge is inherently incomplete, and so we give a chance for new connections to be discovered. Therefore, if a weight survives this regularization by the end of the training (has a non-zero value), its corresponding edge was a contributing interaction to predict the drug response output. To train BDKANN in an end-to-end fashion, we employ the following cost function:

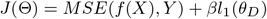

### 3.5 Interpretation

For interpreting the learned BDKANN+ model, we wish to extract a high-attribution sub-network consisting of contributing genes, complexes, and pathways that are highly relevant for predicting the AAC of a given drug. To achieve this goal, we leverage SHAP (SHapley Additive exPlanations) (Lundberg and Lee, 2017), a game theoretic approach for interpreting the predictions of a DNN. SHAP assigns an attribution score to each combination of a DNN node and an input sample, such that a high attribution score indicates that this node plays an important role in the prediction for that sample. We choose to use SHAP, particularly the expected-gradient extension of SHAP (Lundberg, 2018), rather than its alternatives because 1) it unifies various DNN interpretation methods (Alber *et al*., 2019) including LRP (Bach *et al*., 2015) and DeepLIFT (Shrikumar *et al*., 2017), and 2) it combines the idea from integrated gradient to utilize the entire dataset to be used as the background distribution to improve the robustness and consistency of interpretation.

Note that we wish to find a high-attribution sub-network, but the computation of SHAP value only gives the important (high-attribution) nodes within each layer. In order to find such a network, we first compute the SHAP value of each node of each hidden layer and then extract a high-attribution sub-network based on the computed SHAP values in a top-down manner.

The details of the interpretation method can be seen in algorithm 1:

**Algorithm 1.**
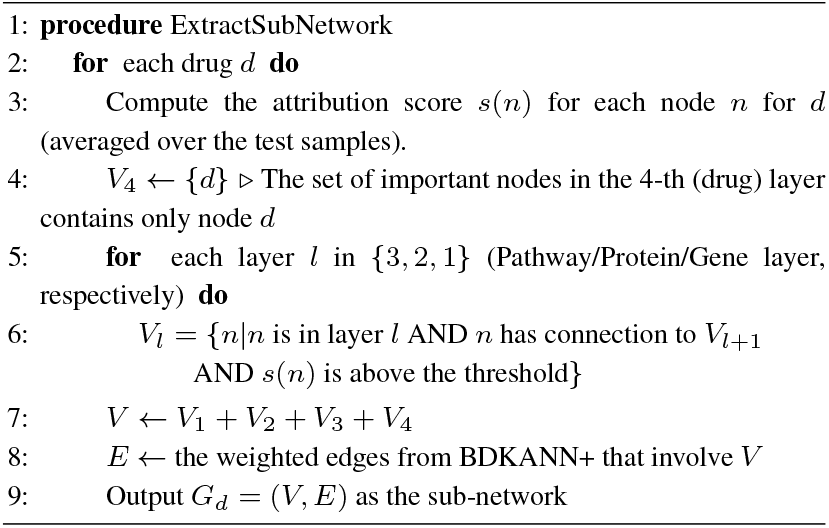
Algorithm for extracting high-attribution sub-networks for each drug.

Given a drug to interpret, we compute the SHAP values for each node in the hidden layer for each sample in the test set. We then take the average over all the samples so that for each drug, we have an attribution score for each node in each hidden layer. We then identify a small set of important nodes in each hidden layer in a top-down manner. More specifically, for the pathway layer, we select a node if 1) it has a connection to important nodes in the upper layer (here it is only the drug node) and 2) its attribution score is above a threshold. The threshold is defined as the relative attribution score ratio, i.e., a node is considered as important if its importance divided by the max importance score in the same layer is above some predefined threshold. After we identify the set of important nodes in the pathway layer, we then find the nodes in the complex layer that are important (according to the threshold) and have connections to the important nodes in the pathway layer. The same process is applied for the complex layer and then the gene layer. We consider the set of important nodes identified for each layer as the nodes of the high-attribution sub-network and annotate the edges of this sub-network with the weight value from the learned BDKANN+ model. By doing this, we extract a high-attribution sub-network, which corresponds to the computation graph that is relevant for predicting the AAC value for a given drug.

## 4 Results

### 4.1 Experimental questions

Our goal is to answer two questions:

1. Does BDKANN outperform knowledge-based and DNN baselines in terms of RMSE?
2. Does the biological knowledge-based structure of BDKANN lead to a more interpretable model and are these interpretations informative?

To answer the first question, we compared our domain knowledge based model with added discovery connections (denoted as BDKANN+) against four baselines. The first baseline is a fully-connected DNN with the number of layers and nodes in each layer tuned as hyper-parameters (denoted as FCv1). This resulted in a network with two hidden layers with 256 and 64 nodes respectively and a total of 172,811 trainable weights. The dimensions of the two hidden layers was intentionally restricted to a max size of 512 neurons in order to keep the size of the FCv1 network comparable to other models used in this domain (Chiu *et al*., 2019). The second baseline (denoted as FCv2) is a fully-connected DNN with the same architecture as the BDKANN+ network (two hidden layers with 4448 and 867 nodes respectively) but without any discovery regularization used. We used dropout for the regularization of these first two baselines. The third baseline, denoted as BDKANN, is a knowledge-based DNN with the same architecture as the BDKANN+ network but only with the connections supported from the domain expert knowledge (illustrated in green in Figure 1). This resulted in quite a sparse network compared to the two fully connected baselines. We did not use dropout for BDKANN and BDKANN+ because we did not want to remove any nodes supported by the domain expert knowledge. Finally, we also compare to a knowledge-based DNN based on GO terms similar to the method of DCELL, which we modified to work for the task of human drug response prediction rather than yeast cell growth (Ma *et al*., 2018). This network took significantly longer to train due to the large size of the network as it was based on the human GO term network.

To answer the second question we calculate the compact hierarchical sub-network for each drug output as explained previously. Then, we investigate the biological entities involved in these sub-networks to see which are shared across drugs and which are drug-specific. Additionally, we can see if we simply recapitulate the known contributors to cancer or if we identify some high-attribution nodes that can point to new cancer drivers or drug targets.

### 4.2 Experimental design

BDKANN+ and all of the baselines were trained and tuned for finding the hyper-parameters that resulted in the lowest RMSE across 10-fold cross validation. Each of the models was then trained and tested on four different data configurations. Models were trained on GDSC and tested on CTRPv2 and conversely trained on CTRPv2 and tested on GDSC. Models were addtionally trained and tested through 10-fold cross-validation on only GDSC data and likewise on only CTRPv2 data. As mentioned previously, there has been significant discussion of the ability to generalize from one dataset to the other despite their similarities, and thus we expect prediction error to be higher in the cross dataset scenario than in the within dataset cross-validation scenario (Consortium *et al*., 2015; Safikhani *et al*., 2016).

The trained BDKANN+ model with discovery regularization (from the data configuration that showed the lowest RMSE) was then passed to the interpretation method as explained above, giving us a high-attribution sub-network for each drug (output).

### 4.3 Prediction results

Figure 2 reports the RMSE of the studied baselines and the two versions of BDKANN on the four different data configurations. As expected, we see that all the models perform better when trained and tested on single dataset rather than training on one dataset and testing on the other. Notably, the FCv1 and BDKANN (with only domain knowledge) perform particularly poorly on the cross-dataset scenarios compared to the other models but the differences between these two models and the rest are less pronounced when trained via cross-validation on one dataset. These two models have significantly fewer parameters (weights) and perhaps fail to learn a strong enough representation of the input to generalize to a different distribution.

**Fig. 2:**
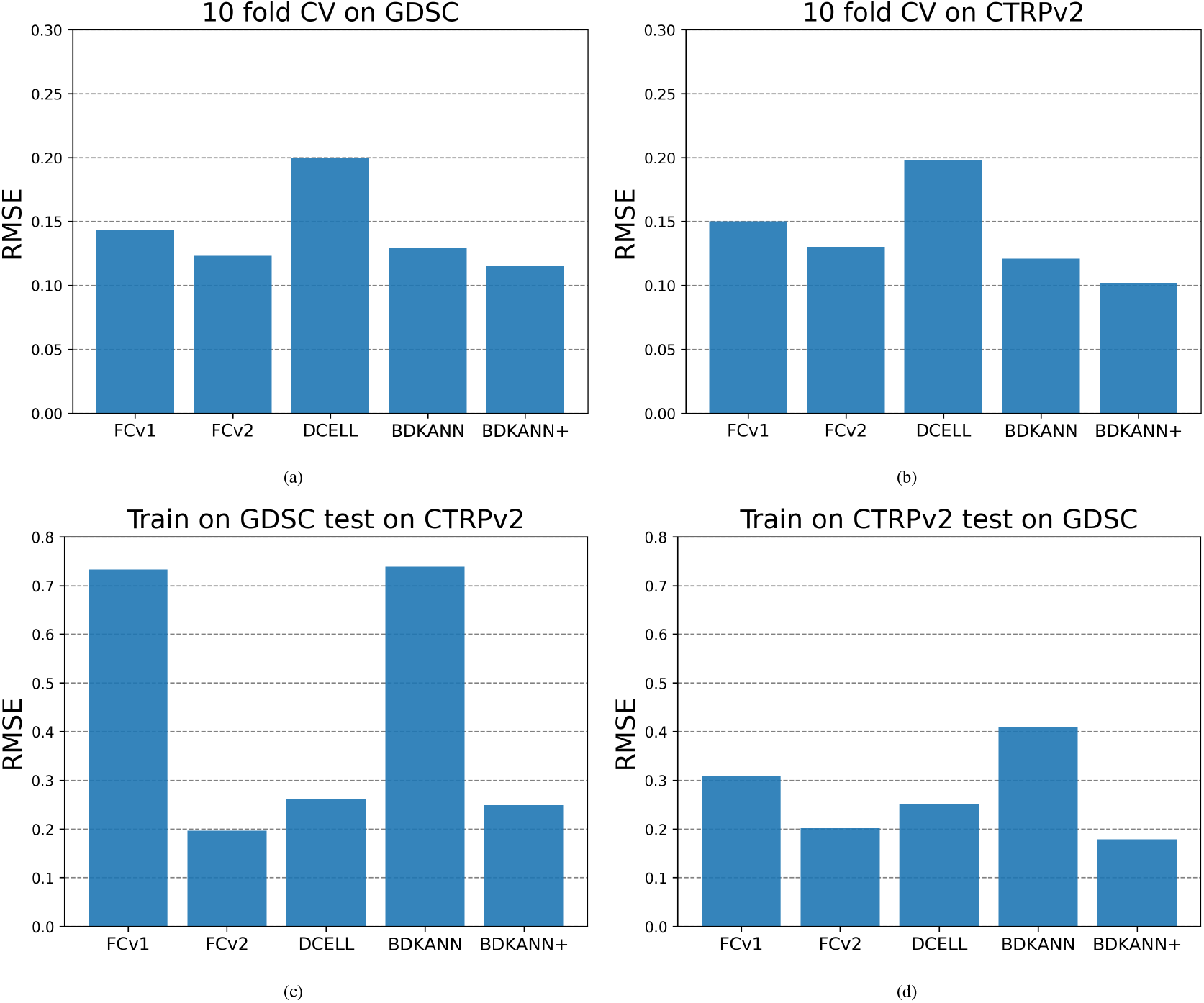
Test RMSE for BDKANN+ and baselines on each of 4 data configurations

Interestingly, the FCv2 model, with the same general architecture as the BDKANN+ model but without any specification of connections, performs well on the cross-dataset scenarios compared to other models and has the lowest RMSE when trained on GDSC and tested on CTRPv2 (Figure 2c). Additionally, the DCELL model also performs relatively well on the cross-dataset scenarios (Figure 2c, 2d) but then performs the worst of all the models when trained and tested on the same dataset (Figure 2a, 2b). It is important to note that the DCELL model tended to predict highly similar values for different inputs, indicating that this model was not able to learn a meaningful function and would probably not generalize well to new data.

Most importantly, the BDKANN+ model outperforms all other baselines on three out of four data configurations and has lower RMSE than both BDKANN and DCELL on all train-test scenarios. The generally high performance of BDKANN+ indicates that the inclusion of biological domain knowledge and the added discovery regularization does not come with the cost of increased prediction error and in fact is an improvement over the other models while also allowing the ability to interpret the model and its predictions.

### 4.4 Interpretation results

The BDKANN+ model, trained on only CTRPv2 data as this gave the lowest RMSE on the test set, was then passed to the interpretation method to produce a high-attribution sub-network. This sub-network tells us which nodes and weights at each layer in the network were important for making the predictions for a given output. Here we show in figures 3a and 3b the extracted high-attribution sub-networks for the cancer drugs Tamoxifen and Selumetinib respectively, which exemplify the different analyses that can be done (high-attribution sub-networks for the remaining 9 drugs were also extracted and included in supplementary material). The IDs at the input layer correspond to Entrez gene names and the IDs at the complex and pathway layer correspond to Reactome identifiers. The thickness of the lines indicate the importance (weight value) of the connection between the nodes.

**Fig. 3:**
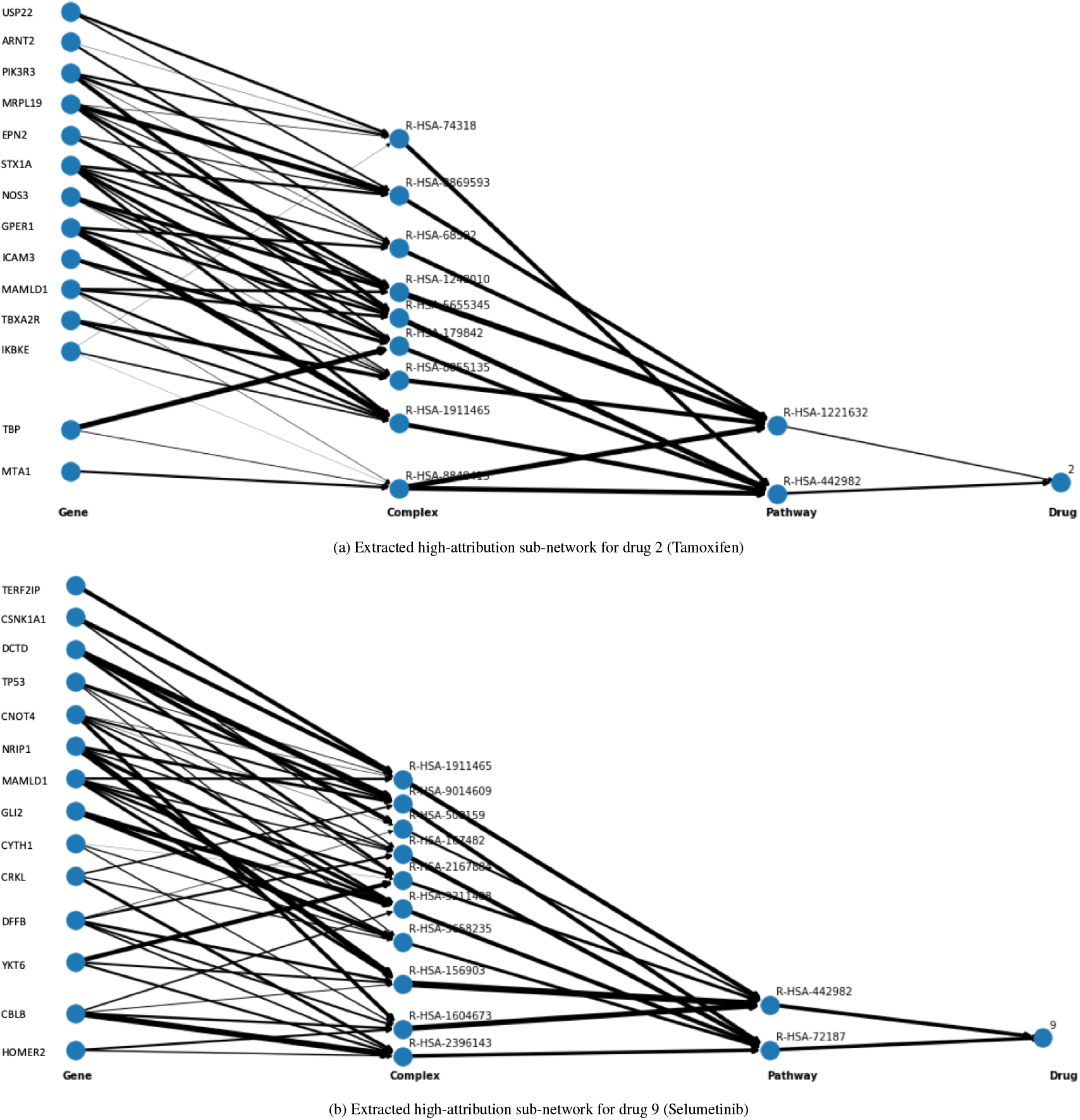
Comparison of high-attribution sub-networks for drugs 2 and 9 (Tamoxifen and Selumetinib). Line thickness indicates weight of the connection.

We can see that there is some overlap between the two high-attribution sub-networks but that they are mostly distinct from each other. At the input layer, the two networks share only the gene mastermind like domain containing 1 (MAMLD1), a gene involved in transcriptional activation and shown to have some involvement in various cancers (Pajtler *et al*., 2017; Qi and Ni, 2019). At the complex layer they share only R-HSA-1911465 which corresponds to the complex of NCOR, HDAC, and TBL1 which are involved in certain signalling cascades for which there is some evidence of involvement in endocrine related cancers Bolós et al. (2007). Lastly, at the pathway layer the two networks share the pathway R-HSA-442982 corresponding to the activation of RAS through calcium influx, RAS being a common oncogene in numerous cancers (Rodenhuis and Slebos, 1992; Viola *et al*., 1986). Thus, the shared entities between the two high-attribution sub-networks appear to have relevance to known mechanisms involved in various cancers.

For a quantitative analysis, table 2 shows the Jaccard similarity for each layer, averaged over all high-attribution sub-networks compared to each other, as well as the similarity between the two high-attribution subnetworks displayed in the text. We can see that the similarity between sub-networks is on average quite low, indicating that each sub-network is distinct from the others. The two high-attribution sub-networks for Tamoxifen and Selumetinib are less similar than average at the gene layer and more similar than average in the complex layer and pathway layer.

**Table 2:** Jaccard similarity coefficient for each layer of high-attribution subnetworks. Row 1: Averaged over all sub-networks with 1 standard deviation shown. Row 2: Similarity between high-attribution sub-networks of drugs 2 and 9 (shown previously).

|  | Gene layer | Complex layer | Pathway layer |
| --- | --- | --- | --- |
| All sub-nets | 0.092 +/- 0.065 | 0.030 +/- 0.044 | 0.090 +/- 0.175 |
| Drugs 2 & 9 | 0.037 | 0.056 | 0.333 |

The overlap between the two sub-networks may suggest some underlying shared phenotype important for the effectiveness of both of these drugs. On the other hand, the fact that the sub-networks are mostly distinct from each other tells us that the BDKANN+ model is finding unique paths in the biological domain knowledge that are useful for predicting each drug output. This is to be expected as Selumetinib and Tamoxifen have distinct biological targets and mechanisms of action and thus they should have distinct sub-networks. However, both drugs have been used to treat the same cancers, including breast cancer, where Selumetinib has shown efficacy in combination with other estrogen-receptor modulators similar to Tamoxifen (Zaman *et al*., 2015). Furthermore, resistance to Selumetinib may be mediated through upregulation of the same RAS pathway identified as common between the two sub-networks (Dry *et al*., 2010). This suggests that the subnetwork both recapitulates the biological domain knowledge and proposes new entities or connections to investigate. We can also see that not all connections in the sub-network are equally important and the layers are not fully connected, showing that a relatively small number of nodes and connections is sufficient for making predictions for each drug.

## 5 Discussion and conclusions

Prediction error results show that BDKANN+ is capable of having comparable or better predictions than baseline methods in a multidrug setting across four different data split scenarios. Importantly, the added discovery regularization does better than only biological domain knowledge (BDKANN) and a fully connected network of the same architecture. Furthermore, the network is interpretable given that its structure is based on biological domain knowledge meaning each node and edge of the network represents a biological entity and the causal relationships that connect them. We show that the extracted high-attribution sub-networks can highlight both general and drug-specific mechanisms and explain which components of the network are important for predicting agivendrug. These sub-networks can be further investigated to identify genes, complexes and pathways that were not previously known to be involved in the action of a drug and possibly discover new drug targets.

Future work will aim to include additional data on more drugs and their pathway connection data as it becomes available, expanding the predictions for more drugs and cell lines. Our general approach of embedding biological domain knowledge in the structure of the model could also be used for tasks beyond drug response prediction such as patient diagnosis or cancer sub-typing. The model could also be expanded to include alternate forms of domain knowledge with different input data, including other omics data types beyond gene expression data. With the increasing need for interpretable models, especially in fields such as cancer drug discovery, our approach serves as a blueprint for designing an accurate predictive model, the decisions of which can also be explained.

## Supporting information

Supplemental figures of high attribution sub networks

