## Supplementary figures and images for "BDKANN - Biological Domain Knowledge-based Artificial Neural Network for drug response prediction"

### Supplemental figures of high attribution sub networks

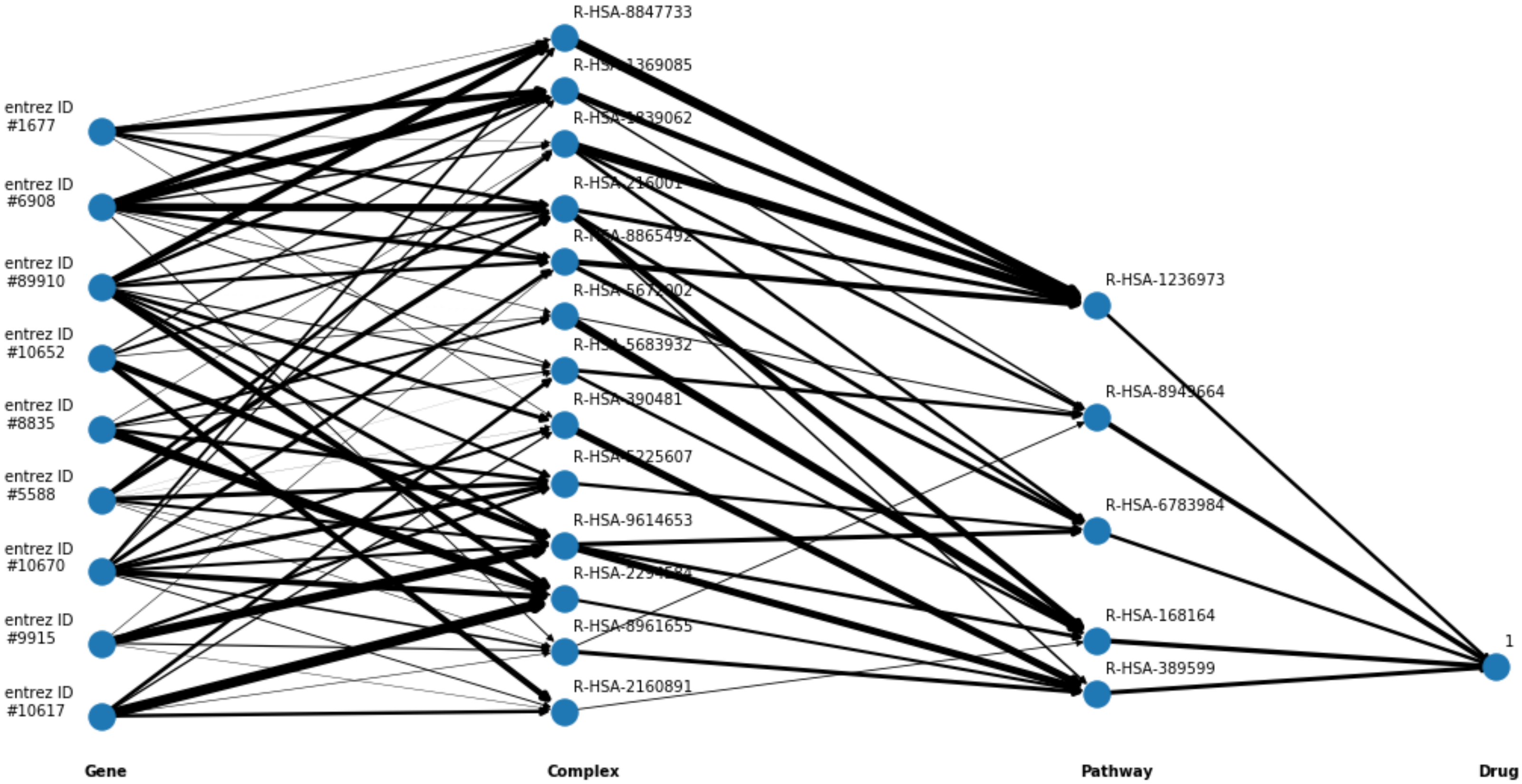

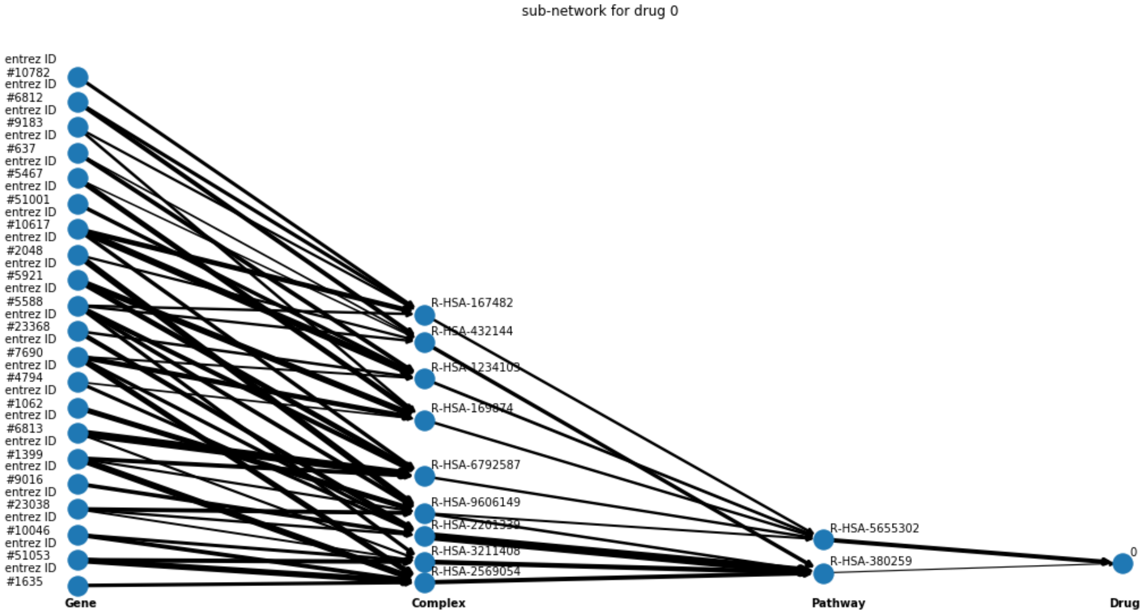

sub-network for drug 3

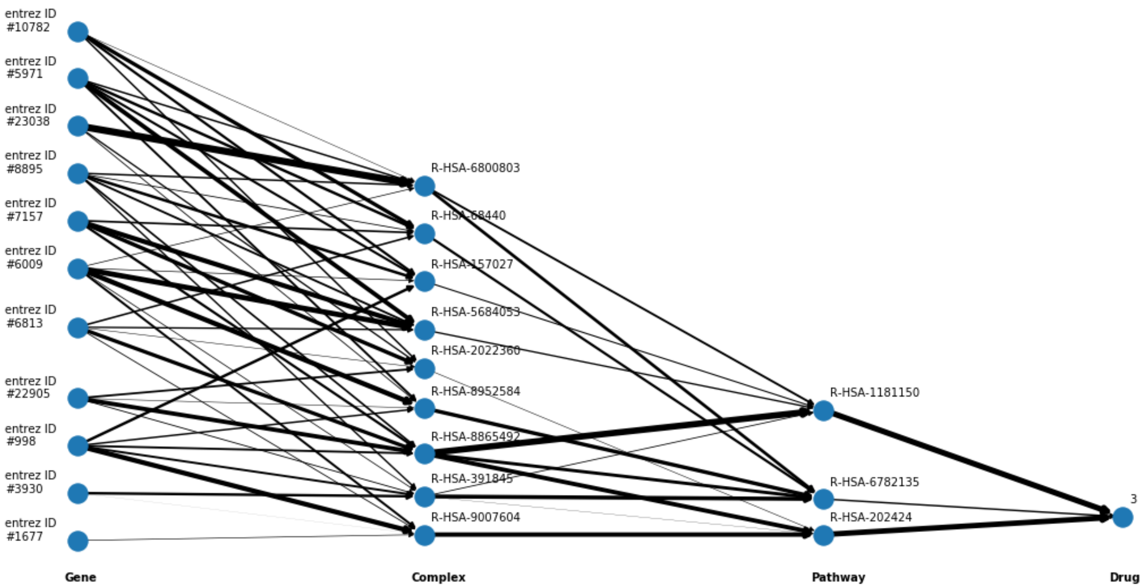

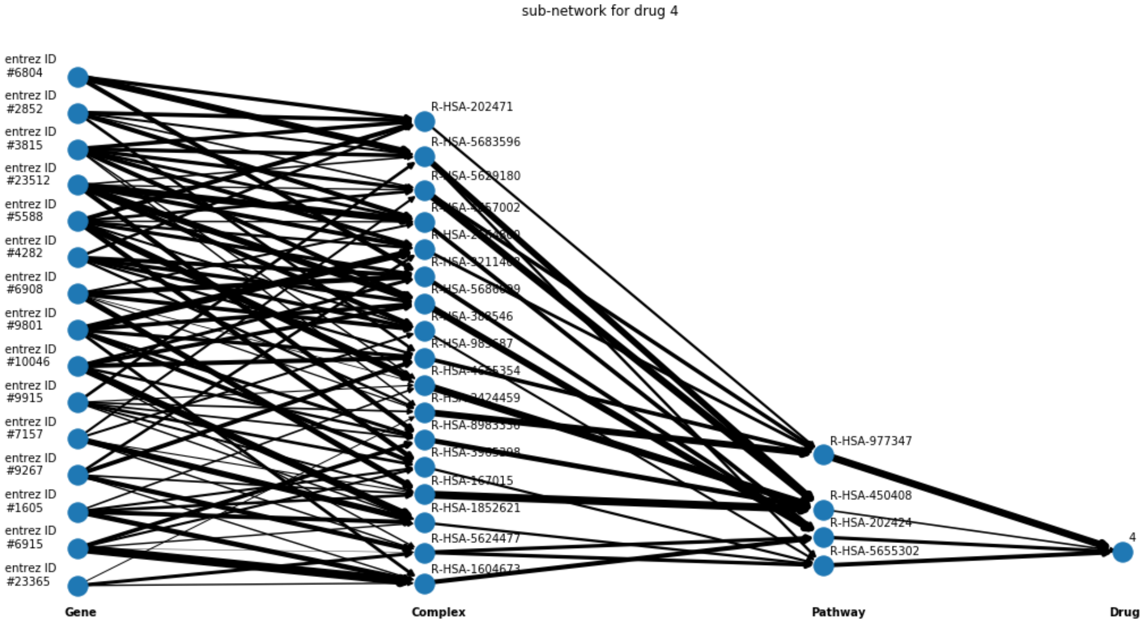

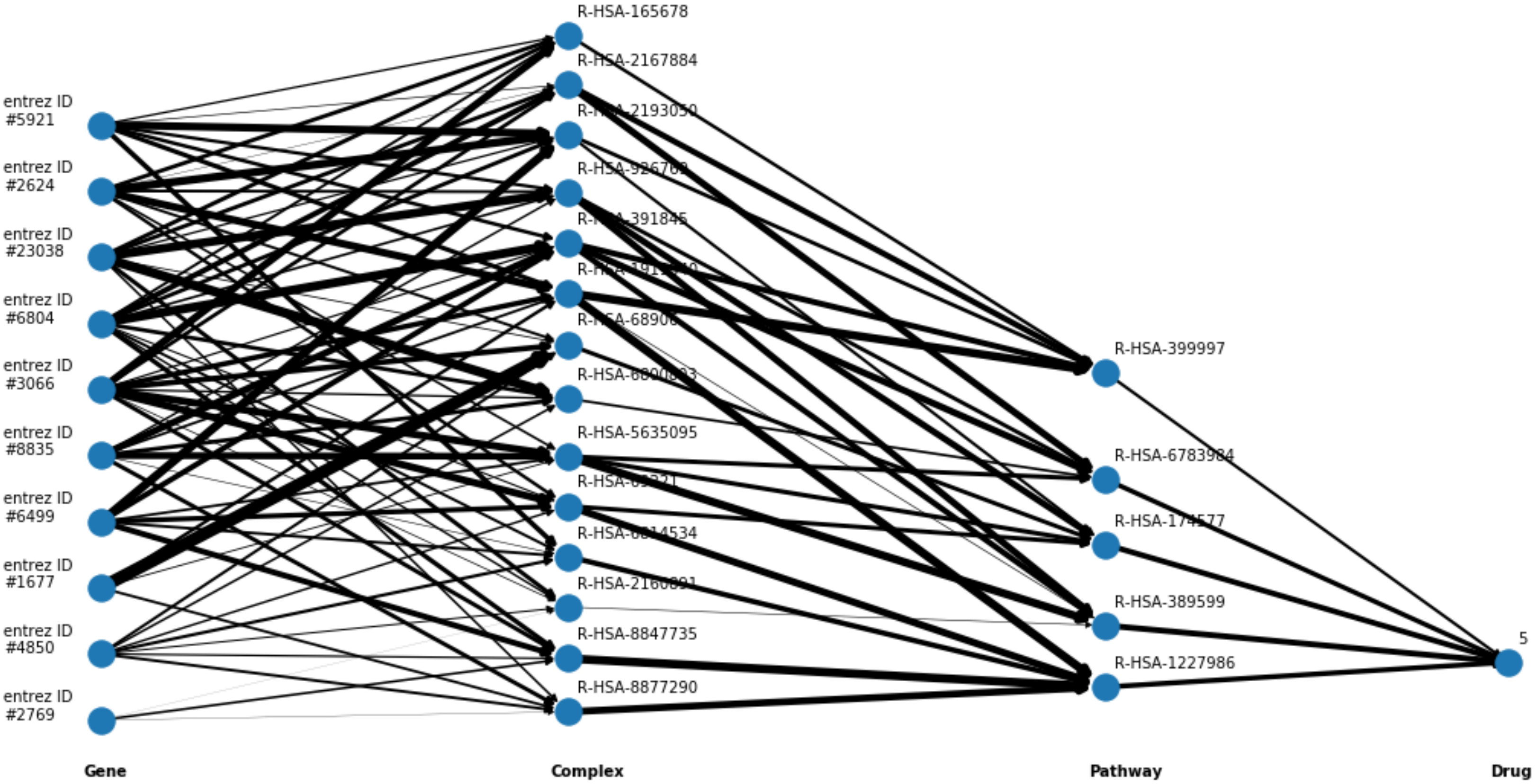

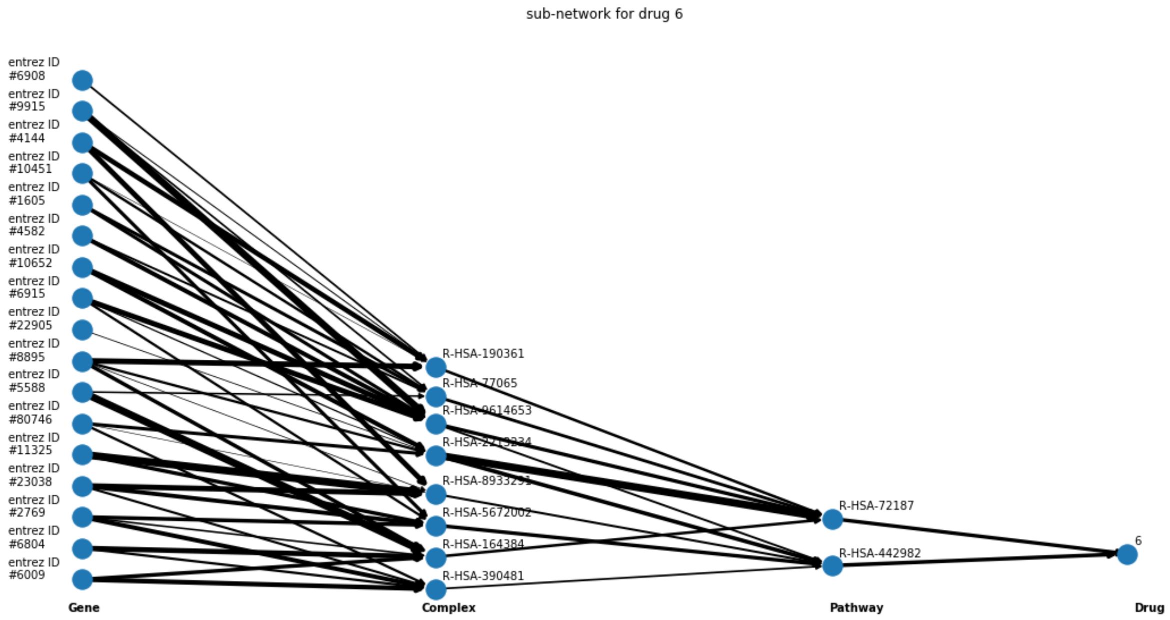

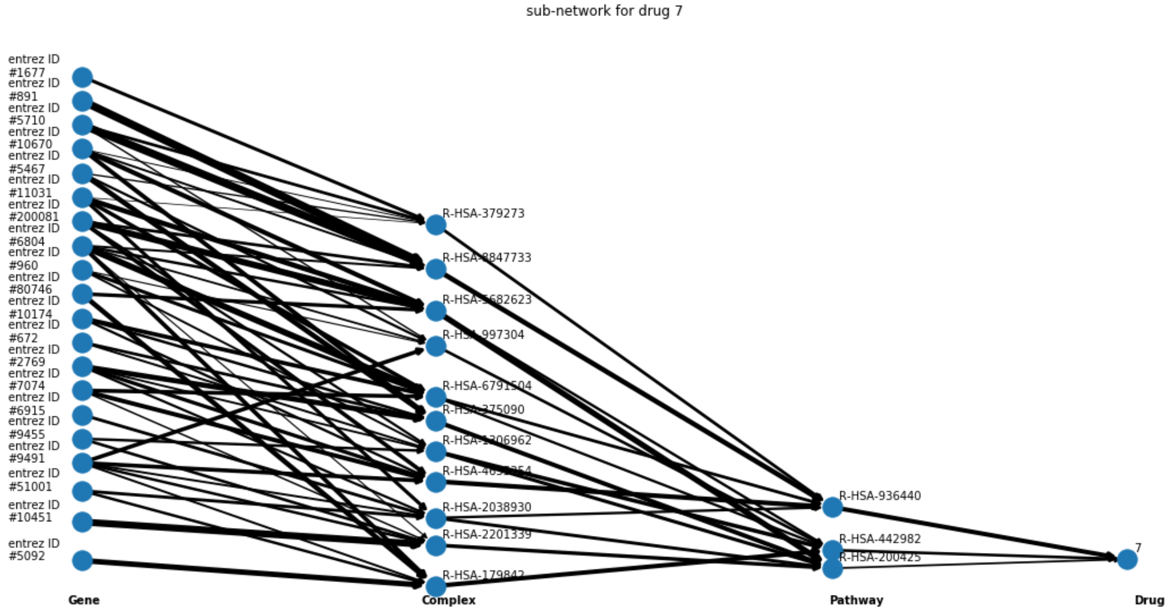

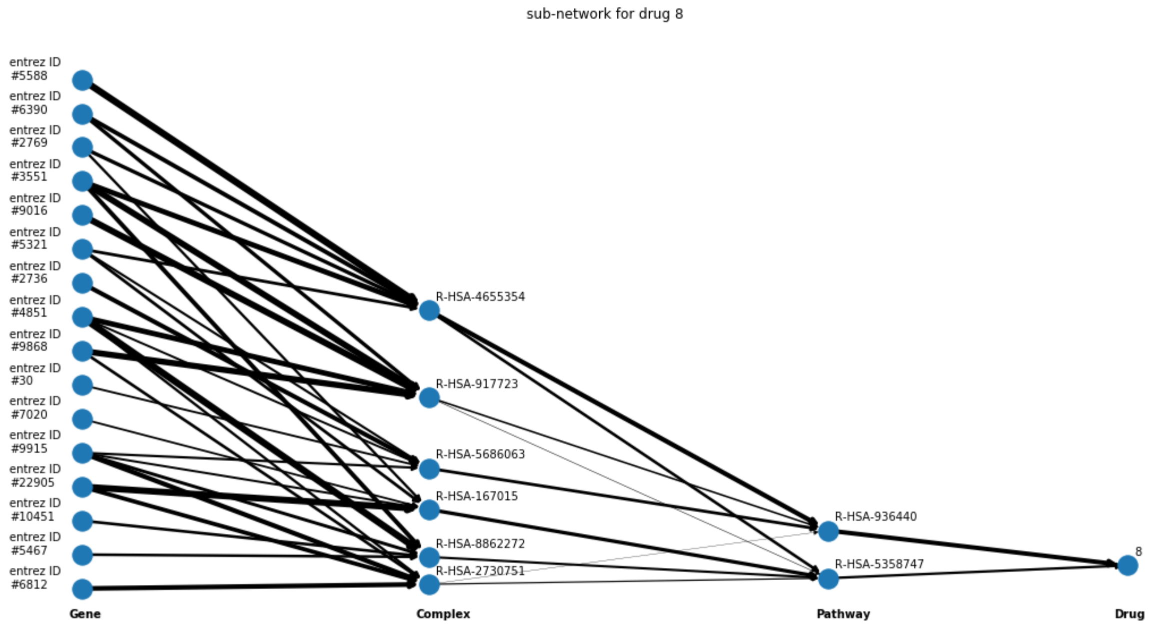

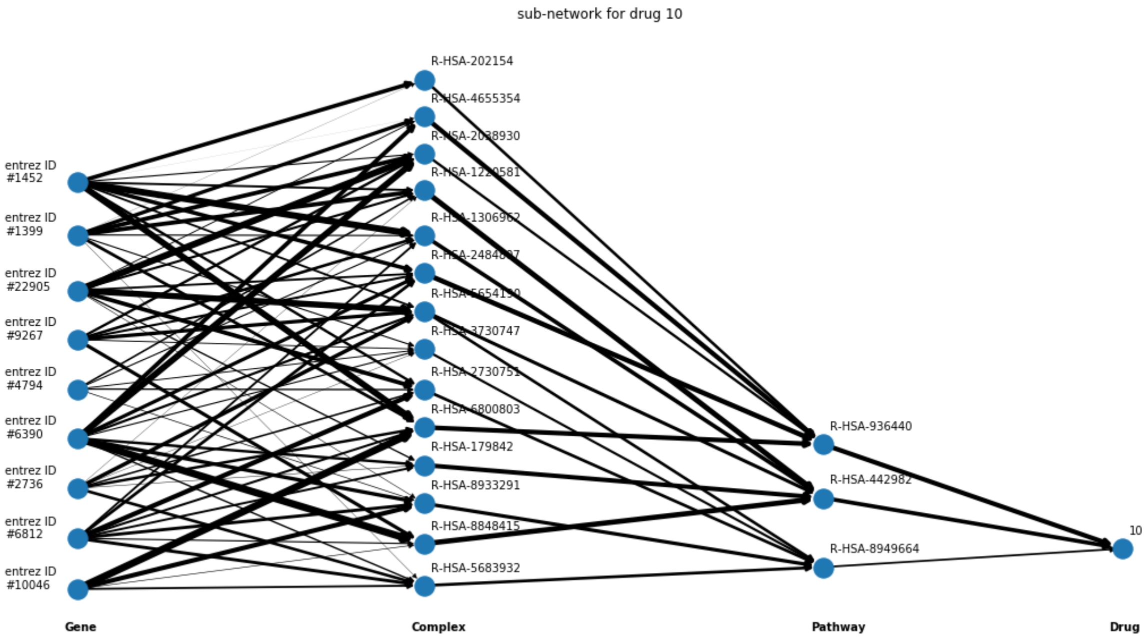
